# The role of the cannabinoid system in fear memory and extinction in male and female mice

**DOI:** 10.1101/2021.06.06.447281

**Authors:** Ikumi Mizuno, Shingo Matsuda, Suguru Tohyama, Akihiro Mizutani

## Abstract

The prevalence of post-traumatic stress disorder (PTSD) is higher in women than in men. Among both humans and mice, females exhibit higher resistance to fear extinction than males, suggesting that differences between sexes in processes of fear extinction are involved in the pathophysiology of such fear-related diseases. Sex differences in molecular mechanisms for fear memory and extinction are unclear. The cannabinoid (CB) system is well known to be involved in fear memory and extinction, but this involvement is based mainly on experiments using male rodents. It has been unclear whether there are sex differences in the role of the CB system in fear memory and extinction. To explore the possibility of such differences, we investigated the effects of pharmacological manipulations of the CB system on the retrieval and extinction of contextual fear memory in male and female mice. WIN55,212-2, a CB receptor (CBR) agonist, augmented the retrieval of fear memory in both sexes, but SR141716 (a CB1R antagonist) did not affect it in either sex. An enhancement of 2-arachidonylglycerol (2-AG, one of the two major endocannabinoids) via JZL184 [an inhibitor of the 2-AG hydrolase monoacylglycerol lipase (MAGL)], augmented the retrieval of fear memory through the activation of CB1R but not CB2R in female mice. In contrast, the enhancement of N-arachidonylethanolamine (AEA, the other major endocannabinoid) via URB597, an inhibitor of an AEA hydrolase (fatty acid amide hydrolase-1) did not show any effects on the retrieval or extinction of fear memory in either sex. WIN55,212-2, SR141716, and JZL184 inhibited fear extinction irrespective of sex. These results suggest that although the role of CB1R in the retrieval and extinction of contextual fear memory is common among males and females, the effects of an increase in the 2-AG level on the retrieval of contextual fear memory differ between the sexes.

## 1. Introduction

The prevalence of post-traumatic stress disorder (PTSD) is higher in women than in men, and the treatment period for PTSD has also been longer in women than in men (Cover et al., 2014). As PTSD patients often show excessive fear memory and an impairment of fear extinction (Milad et al., 2009; Norrholm et al., 2011), it is important to experimentally investigate fear memory and extinction in both sexes in order to understand the mechanisms underlying the sex differences in their prevalence. In both humans and mice, females have exhibited more resistance to fear extinction than males (Kelly and Forsyth, 2007; Matsuda et al., 2015), and female mice tend to exhibit a spontaneous recovery of fear memory compared to males (Matsuda et al., 2015). However, the sex differences in the molecular mechanisms underlying fear memory and extinction are not yet known.

Contextual fear conditioning and extinction comprise one of the most frequently used aspects of behavioral experiments in studies of fear memory and extinction, respectively. Contextual fear conditioning is a type of paired associative learning between a context and an unconditioned stimulus (US) such as a foot shock (Rudy, 1994). After rodents have acquired the associative learning, they begin to exhibit a fear response such as freezing when they are in the conditioned context even without the US. When the rodents are repeatedly presented the context without the US, the fear response decreases. This decrease is called fear extinction, and it is noted that fear extinction does not erase the original fear memory; rather, it forms a new memory of safety that inhibits the expression of the fear response (Herry et al., 2008).

The cannabinoid (CB) receptor (CBR) is one of the molecules related to learning and memory (Marsicano et al., 2002). There are two CBR subtypes: type 1 (CB1R), which is abundantly expressed in the brain; presynapse-resident CB1Rs regulate the release of neurotransmitters (Galiègue et al., 1995; Pan et al., 2009; Yoshida et al., 2011).The other is type 2 (CB2R), which is expressed mainly in immune cells and regulates the expressions of inflammatory cytokines(Carlisle et al., 2002; Galiègue et al., 1995; Tanaka et al., 2020).

The most well-known endocannabinoids (eCBs) are 2-arachidonoyl glycerol (2-AG) and anandamide (AEA), which are degraded by monoacylglycerol lipase (MAGL) and fatty acid amide hydrolases-1 (FAAH), respectively (Blankman et al., 2007; Cravatt et al., 1996; Dinh et al., 2002; Kurahashi et al., 1997). Although 2-AG acts as a full agonist at both subtypes of CBRs (Gonsiorek et al., 2000; Savinainen et al., 2001; Sugiura and Waku, 2000), AEA acts as a partial agonist for CB1R and as a partial agonist or antagonist for CB2R (Glass and Northup, 1999; Gonsiorek et al., 2000; Mackie et al., 1993; Savinainen et al., 2001).

The role of the CB system in fear memory and extinction has been investigated using male rodents. CB1R knockout (KO) male mice and male mice administered the CB1R inverse agonist SR141716 (3 mg/kg) did not exhibit altered retrieval of fear memory (Marsicano et al., 2002), suggesting that the physiological levels of CB1R activation are not involved in the retrieval of fear memory. The effects of CBR agonists and AEA in the retrieval of fear memory are complex and unclear. The CBR agonist WIN55,212-2 (WIN; 2.5 mg/kg) suppressed the retrieval of fear memory in male rodents (Pamplona and Takahashi, 2006), but another CBR agonist called CP55,940 (50 μg/kg) augmented this retrieval in male mice (Llorente-Berzal et al., 2015). A systemic administration of the FAAH inhibitor URB597 (URB; 1.0 mg/kg) suppressed the retrieval of fear memory in male mice (Gobira et al., 2017), but other research groups did not observe such an influence despite the use of the same dose of URB (Llorente-Berzal et al., 2015). The CB2 agonist JWH133 (1 mg/kg) did not affect the retrieval of fear memory in male mice (Llorente-Berzal et al., 2015).

Regarding fear extinction, it was reported that CB1R KO male mice and male mice administered the CB1R inverse agonist SR141716 exhibited resistance to fear extinction (Marsicano et al., 2002). A low dose of WIN (0.25 mg/kg) or the MAGL inhibitor JZL184 (JZL: 0.5 mg/kg) facilitated fear extinction in male rodents (Morena et al., 2018; Pamplona et al., 2006), but a high dose of WIN (2.5 mg/kg) or JZL (8 mg/kg) inhibited it in male rodents (Llorente-Berzal et al., 2015; Pamplona et al., 2006). These findings suggested that the role of CB1R in fear extinction is biphasic in males.

As mentioned above, it is clear that the CB system is involved in fear memory and extinction in male rodents. There have been few investigations of the role of the CB system in fear memory and extinction in female mice. It has thus been unknown whether there are sex differences in these roles of the CB system. Interneurons in the hippocampal area CA1 regulate the retrieval of fear memory (Sans-Dublanc et al., 2020), and sex differences in the effects of URB on the amplitude of an inhibitory post-synaptic current (IPSC) in the hippocampus were observed (Tabatadze et al., 2015). Sex differences in the role of the CB system have also been observed in other behaviors. CB1 KO male mice showed more anxiety-like behavior than wild-type (WT) male mice, but CB1 KO female mice did not show such high anxiety-like behavior (Bowers and Ressler, 2016). It is thus likely that there are also sex differences in the roles of the CB system in fear memory and extinction. To test this hypothesis, we investigated the effects of the pharmacological agents, WIN, SR141716, JZL, and URB (each of which manipulates the CB system) on the retrieval and extinction of contextual fear memory in both male and female mice.

## 2. Materials and methods

### 2.1. Animals

Male and female C57BL/6J mice (14-15 weeks old of age) were purchased from Japan SLC. (Shizuoka, Japan). Mice were housed in open top cage (W220×L320 × H135, Natsume Seisakusho, Tokyo) with 2-4 same sex mice per cage and provided water and food *ad libitum*. Mice were maintained on a 12 h light/dark cycle throughout the study. All behavioral tests were carried out between 9:00 and 17:00. All mice underwent behavioral tests at 15-20 weeks of age. All animal use procedures were approved in advance by the Guidelines for Animal Experimentation of Showa Pharmaceutical University.

### 2.2. Drug treatment

WIN55,212-2 (0.075, 0.75 mg/kg) (Cayman Chemicals, Ann Arbor, MI, USA) was dissolved in saline with 10% dimethyl sulfoxide (DMSO) and 0.1% Tween 20. JZL (4, 8 mg/kg) (Cayman Chemicals) was dissolved in saline with 15% DMSO, 4.25% Tween 20 and 4.25% polyethylene glycol in saline. URB597 (0.3, 1, 3 mg/kg) (Cayman Chemicals) was dissolved in saline with 10% DMSO. SR141716 (1, 5 mg/kg) (Cayman Chemicals) was dissolved in saline with 5% DMSO and 0.1% Tween 20. The CB2R inverse agonist SR144528 (1 mg/kg) (Tocris, Bristol, UK) was dissolved in saline with 6% DMSO and 6% Tween 20. WIN and SR144528 was injected intraperitoneally (i.p.) 30 min before each extinction session or open field test (OFT). JZL was injected i.p. 60 min before each extinction session or OFT. URB was injected i.p. 120 min before each extinction session or OFT. SR141716 was injected i.p. 20 min before each extinction session or OFT. All pharmacological treatments were administered i.p. in an injection volume of 10 ml/kg.

### 2.3. Contextual fear conditioning and extinction

All mice underwent handling (1 min/ day) for 3 consecutive days until 2 days before the behavior tests. Contextual fear conditioning and extinction were performed according to our previous study (Matsuda et al., 2018).

On day 1, the mouse was placed in a conditioning chamber (22.8×19.7×13 cm) for 20 min as habituation. On day 2, the mouse received three foot-shocks (2 s, 0.75 mA, 60-120 s intertrial interval) after a 180-s acclimation period in the conditioning chamber. The mouse was subsequently returned to its home cage after a last 180-s foot-shock. We used the data of the average of the percentage freezing during the last 2 min in fear conditioning as the POST value. After matching for equivalent levels of freezing in POST, conditioned mice were divided into groups of two or four mice for the injection of vehicle or drugs.

On day 3, the mice were re-exposed to the conditioning chamber for 6 min (TEST) for the investigation of the effects of the drugs on only the retrieval of fear memory or for 20 min (FE1) for the investigation of the effects of the drugs on both the retrieval of and extinction of fear memory. On days 4-7 (FE2-FE5), the latter group of mice was re-exposed to the conditioning chamber for 20 min. On day 28 [recall of fear extinction (RE)], the mice were re-exposed to the conditioning chamber for 6 min. Each chamber was cleaned with 70% ethanol before each session. The behavior of each mouse was monitored and recorded inside the chamber by a CDD camera mounted on the top of the sound-attenuating box. We defined immobility (freezing) as an immobility duration of 1 second. Freezing (no visible movement except for respiration) was scored by the automatic program TimeFZ4 (O’Hara & Co., Wilmette, IL) and then converted into a percentage.

### 2.4 Open field test

The open field apparatus was a square field (W50 × L50 × H30 cm) made of white acrylic material. The mouse was placed in the corner of apparatus at the beginning of the test and allowed to move freely for 20 min. The total distance and total center time were recorded. The center area was 16 ×16 cm. The total distance and the total center time were evaluated as an index of locomotor activity and anxiety, respectively. The data analysis was performed using TopScan (Clever Sys, Reston, VA).

### 2.5 Determination of estrous cycle

We collected vaginal smears from the female mice between 08:00 and 10:00 according to our previous study (Matsuda et al., 2018). To investigate the effects of the estrous cycle on the TEST-day values, FE, and OFT-day values, we examined one vaginal smear per mouse per day for 5 days (FE1 through FE5) or from 2 days before the TEST or OFT day to 2 days after the TEST or OFT day. We calculated the percentage of mice in the proestrus phase P(%) in each group by the day as follows: P(%) = [number of mice in the proestrus/(total number of mice in the proestrus, estrus, and diestrus)] × 100.

### 2.6 Statistical analysis

We used the data from the average of percentage freezing during the initial 6 min on day 3 and on day 28 as TEST and RE, respectively. Further, we used the data from the average of percentage freezing at 2-min intervals in each extinction session to measure and calculate fear extinction. For fear conditioning and extinction, we used one-way (group) and multiple (bin, day and group) analysis of variance (ANOVA) to detect significant differences, respectively. In the OFT we used the data from the average of moving distance at 2-min intervals and the total duration time in the center area to measure the total distance and center time, respectively. For the center time and total distance, we used a one-way (group) ANOVA or Student’s t-test and a two-way (time and group) repeated ANOVA, respectively, for significant differences. We used a post-hoc Bonferroni test or Dunnett’s test for multiple comparisons. For all analyses, the level of statistical significance was set at p<0.05. All statistical were performed using SPSS 22.0 J for Windows (SPSS Japan, Tokyo). The data are presented at mean ± SEM.

## 3. Results

The estrous cycle data of the female mice in each experiment are illustrated in Supplemental Figures S1–S3, which demonstrate that the percentage of mice in the proestrus phase were comparable between the drug-administered groups in all experiments.

### 3. 1. Effect of WN55,212-2 on the retrieval and extinction of fear memory

To investigate the role of the activation of CB1R on the retrieval and extinction of fear memory in males and female mice, we administered WIN before each contextual fear extinction session.

In TEST, there were significant differences in %freezing among groups in the male mice (main effect of group: *F*_(2, 33)_ = 10.709, *p*<0.001) and the female mice (main effect of group; *F*_(2, 30)_ = 6.054, *p*< 0.01). The post-hoc analysis revealed that the mice treated with WIN (0.75 mg/kg) showed a significantly higher %freezing compared to the mice treated with the vehicle-alone among both the males (*p*< 0.001; Fig. 1B) and females (*p*< 0.01; Fig. 1E). These results suggested that WIN augmented the retrieval of fear memory in both sexes.

**Fig. 1.**
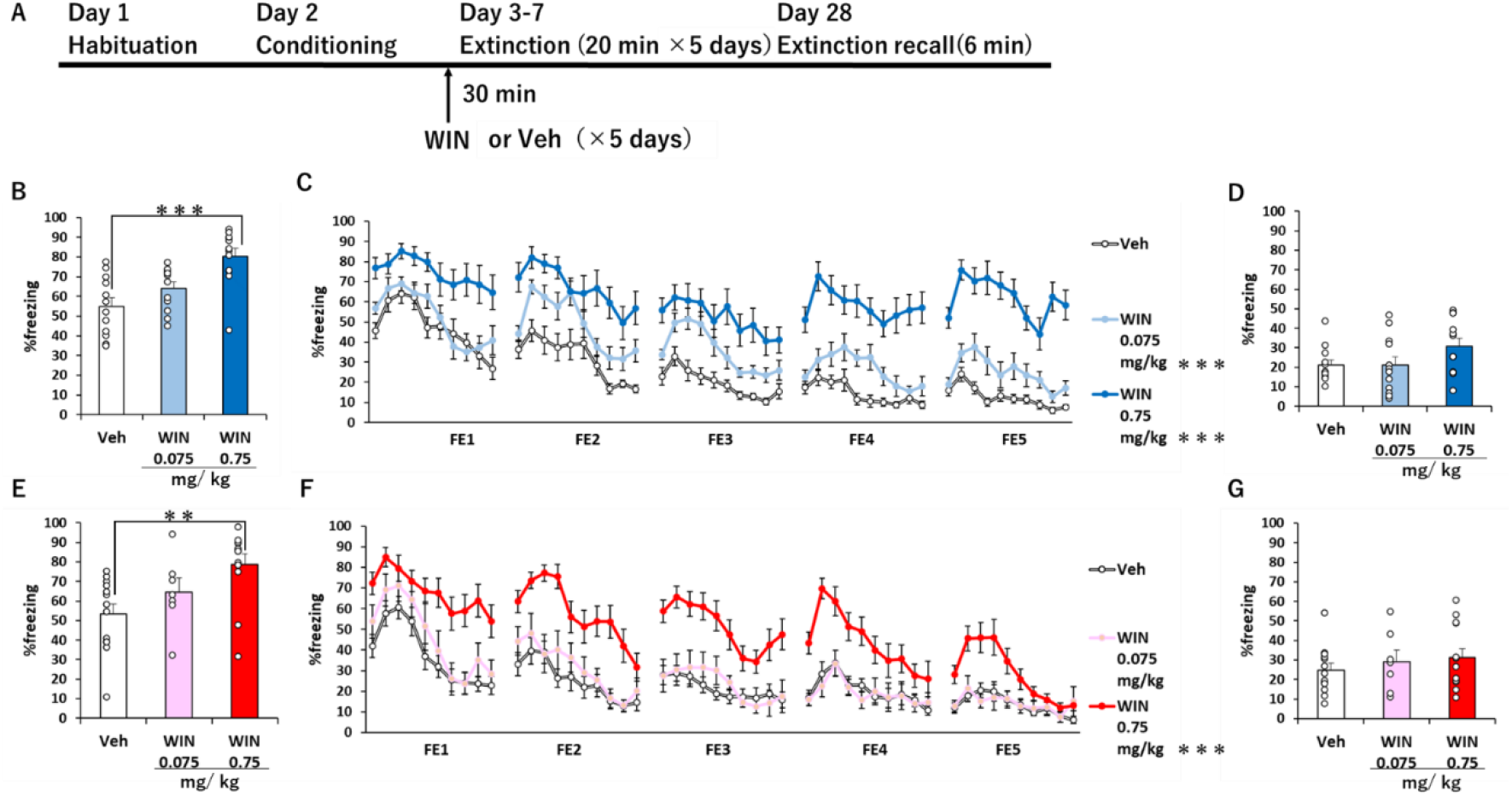
WIN augmented the retrieval of fear memory and suppressed fear extinction in both sexes. A: The experiment 1 procedure. The *arrow* indicates the time point of drug injection performed 30 min before each FE. The percentage of freezing during the TEST, FE, and RE in males (**B–D**) and females (**E–G**), respectively. *White circles* represent each animal’s data (n=7– 13/ group). \*\**p*< 0.01, \*\*\**p*< 0.001, WIN (0.75 mg/kg and 0.075 mg/kg) vs. Veh.

Regarding fear extinction, all groups showed significant differences in %freezing among FE in the males (main effect of FE day; *F*_(4, 165)_ = 19.391, *p*< 0.001) and females (main effect of FE day; *F*_(4, 150)_ = 20.016, *p*< 0.001). The post-hoc analysis revealed that all groups showed significantly lower %freezing on day FE5 compared to day FE1 (*p*< 0.001). These results indicated that all groups acquired fear extinction. The %freezing during the extinction sessions differed among groups in the males (main effect of group; *F*_(2, 165)_ = 19.391, *p*< 0.001; interaction of bin × group; *F*_(12.084, 996.905)_ = 1.859, *p*< 0.05) and the females (main effect of group; *F*_(2, 150)_ = 58.665, *p*< 0.001; interaction of bin × group × FE day; *F*_(51.245, 960.841)_ = 1.592, *p*< 0.001). The post-hoc analysis revealed that the males treated with WIN (0.075 mg/kg and 0.75 mg/kg) showed significantly higher %freezing than the males treated with vehicle (*p*< 0.001; Fig. 1C, Suppl. Fig. S4). Moreover, the females treated with WIN (0.75 mg/kg) showed significantly higher %freezing than the females treated with vehicle (*p*< 0.001; Fig. 1F, Suppl. Fig. S4). However, such differences disappeared in RE in the males (*F*_(2, 33)_ = 2.260, *p*= 0.120; Fig. 1D) and in the females (*F*_(2,30)_ = 0.607, *p*=0.551; Fig. 1G). These results suggested that WIN inhibited fear extinction in both sexes but did not affect the retention of fear extinction in either sex.

### 3.2 Effect of SR141716 on the retrieval and extinction of fear memory

Investigations of male rodents showed that the 2-AG and AEA levels in fear-related brain regions such as the basolateral amygdala (BLA) and hippocampal area CA1 increased after the retrieval of a fear memory (Marsicano et al., 2002; Segev et al., 2018). Although the eCB levels were not directly measured in those studies, the levels of eCBs may increase during the TEST period because eCBs are synthesized on demand by depolarizing stimuli (Di et al., 2005; Jung et al., 2005; Stella and Piomelli, 2001). We therefore investigated the role of physiological levels of eCBs in the retrieval and extinction of fear memory by using SR141716, a CB1R antagonist. In the TEST data, there were no significant differences in %freezing among groups in the males (*F*_(2, 33)_ = 0.129, *p*= 0.880; Fig. 2B) or females (*F*_(2, 31)_ = 0.396, *p*= 0.676; Fig. 2E).

**Fig. 2.**
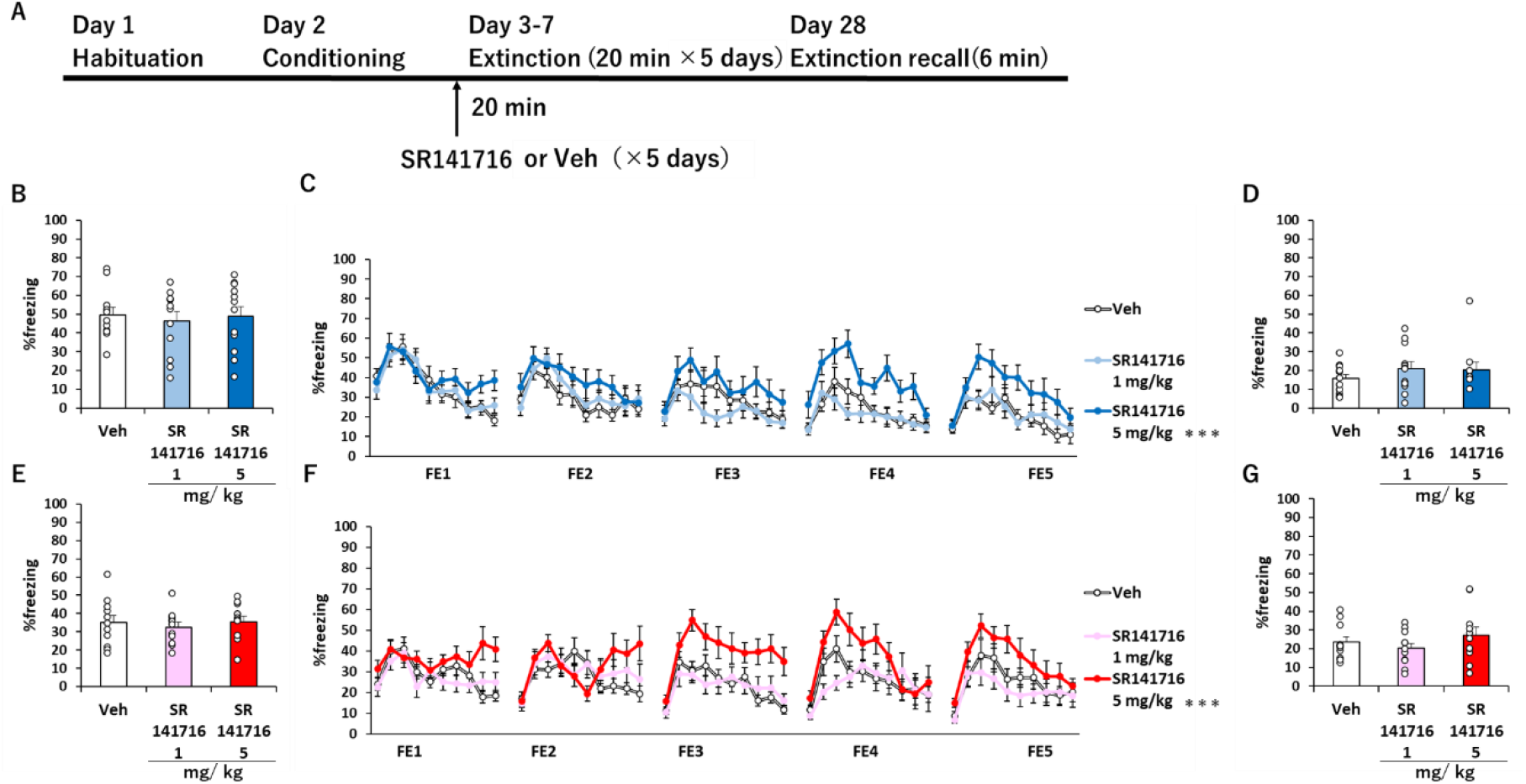
SR141716 did not affect the retrieval of fear memory in either sex but suppressed fear extinction in both sexes. **A:** The procedure of experiment 2. The *arrow* indicates the time point of drug injection performed 20 min before each FE. The percentage of freezing during the TEST, FE, and RE in the males (**B–D**) and females (**E–G**), respectively. *White circles*: each animal’s data (n = 11–12/ group). \*\*\**p* < 0.001 SR141716 (5 mg/kg) vs. Veh.

Regarding fear extinction, all groups showed differences in %freezing among FE days in both the males (main effect of FE day: *F*_(4, 160)_ = 4.849, *p*< 0.01; Fig. 2C) and females (interaction of bin × group × FE day; *F*_(57.031, 1069.340)_ = 1.817, *p*< 0.001; Fig. 2F). The post-hoc analysis revealed that all groups in males showed significantly lower %freezing on day FE5 compared to day FE1(*p*< 0.01), and all groups of females showed significantly lower %freezing in the first bin of day FE5 compared to day FE1 (*p*< 0.01). These results indicated that all groups acquired fear extinction.

On the other hand, the %freezing during the extinction sessions differed among groups in both the males (main effect of group; *F*_(2, 160)_ = 12.091, *p*< 0.001; Fig. 2C) and females (main effect of group; *F*_(2, 150)_ = 14.577, *p*< 0.001; interaction of bin × group × FE day; *F*_(57.031, 1069.340)_ = 1.817, *p*< 0.001; Fig. 2F). The post-hoc analysis revealed that the mice treated with SR141716 (5 mg/kg) showed significantly higher %freezing than the mice treated with vehicle in both the males (*p*< 0.001; Fig. 2C) and females (*p*< 0.001; Fig. 2F, Suppl. Fig. S5). However, such differences disappeared in RE in both the males (*F*_(2, 33)_ = 0.733, *p*= 0.488; Fig. 2D) and the females (*F*_(2,31)_ = 1.011, *p*= 0.376; Fig. 2G). These results indicated that SR141716 inhibited fear extinction in both sexes but did not affect the retention of fear extinction in either sex.

### 3.3 Effect of JZL on the retrieval and extinction of fear memory

Next, to investigate the role of excess levels of 2-AG in the retrieval and extinction of fear memory, we administered JZL before each extinction session. The TEST results revealed that while there was no significant difference in %freezing among groups in the males (main effect of group; *F*_(2, 33)_ = 0.012, *p*= 0.988; Fig. 3B), there was a significant difference in the females (main effect of group; *F*_(2, 56)_ = 4.218, *p*< 0.05; Fig. 3E). The post-hoc analysis revealed that the female mice treated with JZL (8 mg/kg) showed significantly higher %freezing compared to the females treated with vehicle (*p*< 0.05; Fig. 3E). These results indicated that excess levels of 2-AG augmented the retrieval of fear memory in females but not in males.

**Fig. 3.**
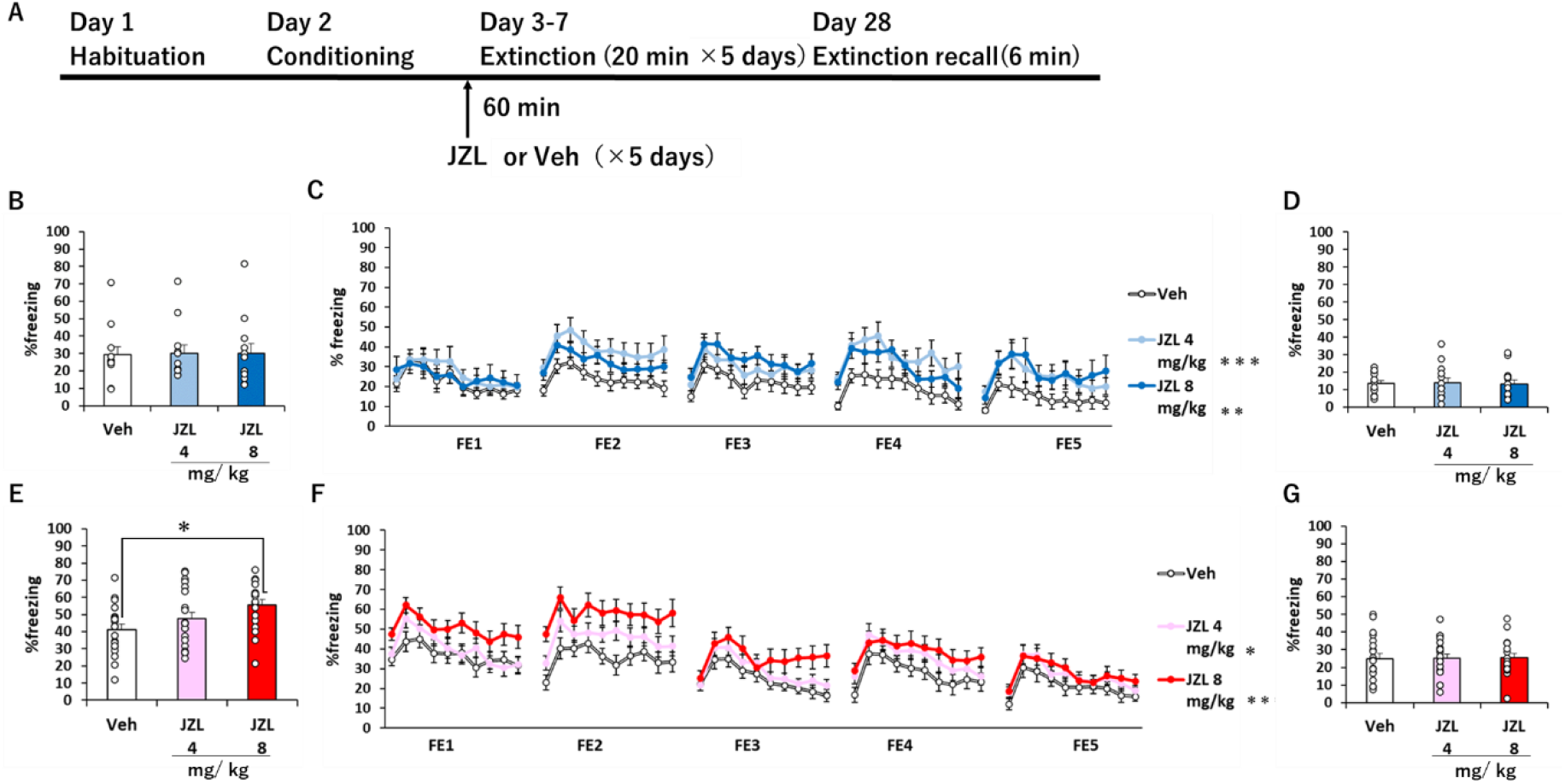
JZL augmented the retrieval of fear memory in only females but suppressed fear extinction in both sexes. **A:** The procedure of experiment 3. The *arrow* indicates the time point of drug injection that was performed 60 min before each FE. The percentage of freezing during the TEST, FE and RE in males (**B–D**) and females (**E–G**), respectively. *White circles*: each animal’s data (n = 12–21/ group). \**p*< 0.05, \*\**p*< 0.01 and \*\*\**p*< 0.001, JZL (4 mg/kg and 8 mg/kg) vs. Veh.

In the fear extinction protocol, all groups showed significant differences in %freezing among FE days in both the males (interaction of bin × FE day; *F*_(26.360, 1087.365)_ = 1.837, *p*<0.01; Fig. 3C) and females (main effect of FE day; *F*_(4, 280)_ = 20.161, *p*< 0.001; Fig. 3F). The post-hoc analysis revealed that all groups of males showed significantly lower %freezing in the first bin of day FE5 compared to that of day FE1 (*p*< 0.01), and all groups of females showed significantly lower %freezing on day FE5 than day FE1(*p*< 0.001). Notably, the %freezing during the extinction sessions differed among groups in both the males (main effect of group; *F*_(2, 165)_ = 8.859, *p*< 0.001; Fig. 3C) and females (main effect of group; *F*_(2, 280)_ = 17.408, *p*< 0.001; Fig. 3F). The post-hoc analysis revealed that the mice treated with either the low dose or the high dose of JZL showed significantly higher %freezing than the mice treated with vehicle among both the males (*p*< 0.001, *p*< 0.01, respectively; Fig. 3C) and females (*p*< 0.05, *p*< 0.01, respectively; Fig. 3F). However, such differences disappeared in RE in both the males (*F*_(2, 34)_ = 0.157, *p*= 0.855; Fig. 3D) and females (*F*_(2,57)_ = 0.020, *p*= 0.980; Fig. 3G). These results indicated that excess levels of 2-AG inhibited fear extinction in both sexes but did not affect the retention of fear extinction in either sex.

### 3.4 JZL augmented fear memory through the activation of CB1R, but not CB2R, in the females

Since 2-AG binds to both CB1R and CB2R, we sought to determine which receptor is involved in the augmentation of retrieval of fear memory in female mice by a co-administration of a high dose of JZL with the respective selective antagonists of CB1R and CB2R. When SR141716 was co-administered with JZL, the %freezing in the TEST period differed significantly among the groups of female mice (main effect of group; *F*_(3, 58)_ = 3.039, *p*< 0.05; Fig. 4B). The post-hoc analysis revealed that the administration of JZL+vehicle tended to increase %freezing compared to only vehicle (*p*= 0.076, Fig 4B). However, such an increase was reduced by SR141716.

**Fig. 4.**
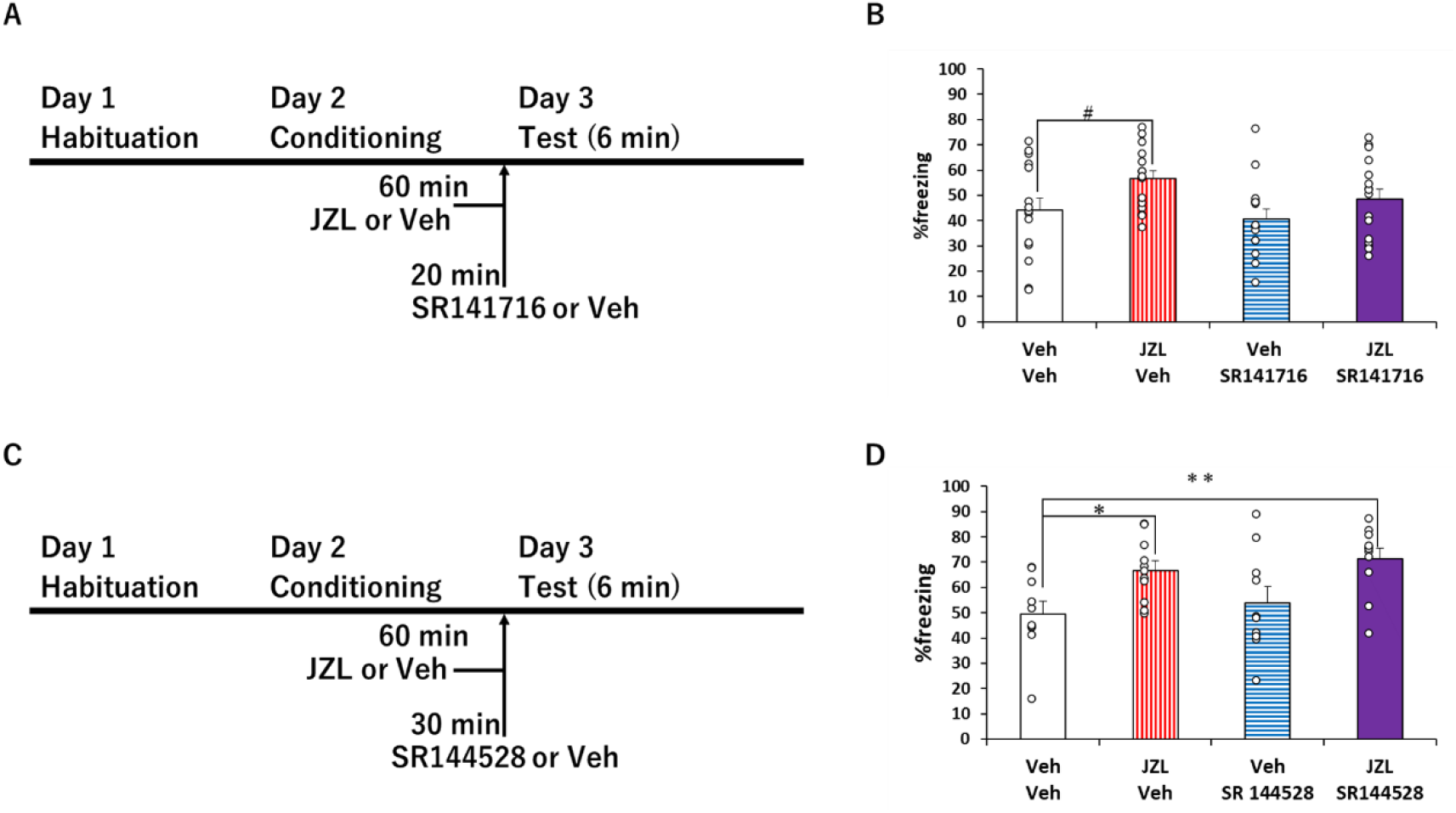
Systemic administration of SR141716, but not SR144528, prevented the augmentation of retrieval of fear memory by JZL. Experiment 4, in which JZL and SR141716 (**A**) or SR144528 (**B**) were injected before TEST. The *arrow* indicates the time point of drug injection when JZL or Veh and SR141716 or Veh was injected 60 min and 20 min, respectively, or JZL or Veh and SR144528 or Veh was injected 60 min and 30 min, respectively, before TEST. The percentage of freezing during the TEST in which JZL and SR141716 (**C**) or SR144528 (**D**) were injected before TEST. *White circles*: each animal’s data (n = 10-16/ group). #: *p*=0.076 JZL (8 mg/kg) vs. Veh. \**p*< 0.05, \*\**p*< 0.01, JZL (8 mg/kg) and JZL (8 mg/kg) + SR144528 (1 mg/kg) vs. Veh.

We next combined JZL and the CB2 antagonist SR144528, and the results demonstrated that the %freezing in the TEST period differed significantly among the groups of female mice (main effect of group; *F*_(3, 38)_ = 4.677, *p*< 0.01; Fig. 4D). The post-hoc analysis revealed that not only administration of JZL+vehicle but also the administration of JZL and SR144528 significantly increased the %freezing compared to only vehicle (*p*< 0.05, *p*< 0.01, respectively; Fig. 4D). Thus, 2-AG augmented the retrieval of fear memory through the activation of CB1R, but not CB2R, in female mice.

### 3.5 Effect of URB on the retrieval and extinction of fear memory

To investigate the roles of excess levels of AEA in the retrieval and extinction of fear memory, we administered URB before each extinction session. In the TEST period, while there no significant difference in the %freezing among groups in the males (main effect of group; *F*_(3, 45)_ = 0.726, *p*= 0.542; Fig. 5B), there was a significant difference in %freezing among groups in the females (main effect of group; *F*_(3, 44)_ = 2.933, *p*< 0.05; Fig. 5E). The post-hoc analysis revealed that among the female mice, those treated with URB (3 mg/kg) tended to show higher %freezing than those treated with vehicle alone (*p*= 0.057).

**Fig. 5.**
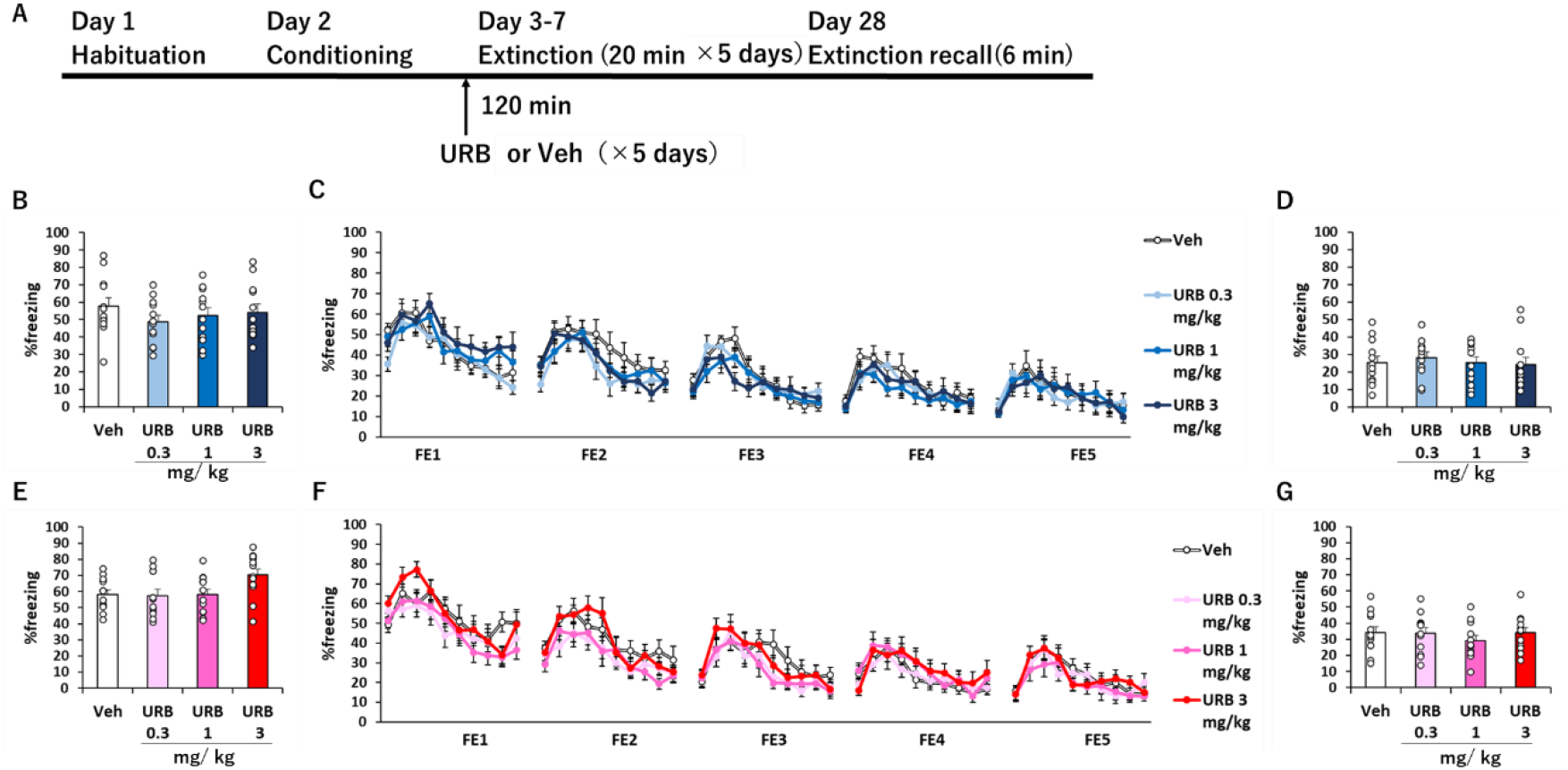
URB did not affect the retrieval or extinction of fear memory in either sex. A: The procedure of experiment 5. The *arrow* indicates the time point point of drug injection that was performed 120 min before each FE. The percentage of freezing during the TEST, FE, and RE in males (**B–D**) and females (**E–G**), respectively. White circles: each animal’s data (n= 12/ group).

Since this result was only a tendency, we performed an additional experiment to investigate the effect of URB (3 mg/kg) on the retrieval of fear memory. There was no significant difference in %freezing between the treated group of either sex (Suppl. Fig. S6). These results indicated that excess levels of AEA did not affect the retrieval of fear memory in either sex. In fear extinction, all groups showed significant differences in %freezing among FE in both the males (main effect of group; *F*_(4, 220)_ = 27.609, *p*< 0.001; Fig. 5C) and females (main effect of group; *F*_(4, 220)_ = 49.636, *p*< 0.001; Fig. 5F). The post-hoc analysis revealed that all groups showed significantly lower %freezing on day FE5 compared to day FE1 (*p*< 0.001). These results indicated that all groups acquired fear extinction.

Although there was no significant fear-extinction difference in the %freezing among groups in the males (main effect of group; *F*_(3, 220)_ = 0.714, *p*= 0.545; interaction of bin × Group; *F*_(83.994, 370332.197)_ = 0.861, *p*= 0.290; interaction of bin × Group × FE day; *F*_(20.998, 370332.197)_ = 1.147, *p*= 0.809; Fig. 5C), there was a significant difference in %freezing among groups in the females (main effect of group; *F*_(3, 220)_ = 3.212, *p*< 0.05; Fig. 5F). However, the post-hoc analysis revealed no significant difference in %freezing among groups in the females, and there was no significant difference in %freezing during RE in the males (*F*_(3, 45)_ = 0.230, *p*= 0.875; Fig. D) or the females (*F*_(3, 45)_ = 0.528, *p*= 0.665; Fig. 5G). These results indicated that excess levels of AEA did not affect fear extinction or the retention of fear extinction in either sex.

### 3.6 Effects of drugs targeting the CB system on locomotor activity or anxiety-like behavior

Because we defined the %freezing as no visible movement except respiration in the mice, it is possible that the alterations in %freezing were a result of the effect of the drugs administered on locomotor activity. To examine this possibility, we investigated the effects of the drugs that altered %freezing in the present study on locomotor activity by conducting the open field test (OFT).

The results of the ANOVA and Student’s t-test in each experiment are summarized in Supplemental Table S1, demonstrating that none of the drugs used affected the total distance or the duration of the time spent in the center of the chamber in both the males and females (Fig. 6); WIN, JZL, and SR141716 thus did not appear to affect the locomotor activity or anxiety-like behavior of the mice. The alterations in % freezing during the TEST and extinction session by the drugs targeting the CB system in this study therefore represent the respective drugs’ effects on the retrieval and extinction of fear memory in the mice.

**Fig. 6.**
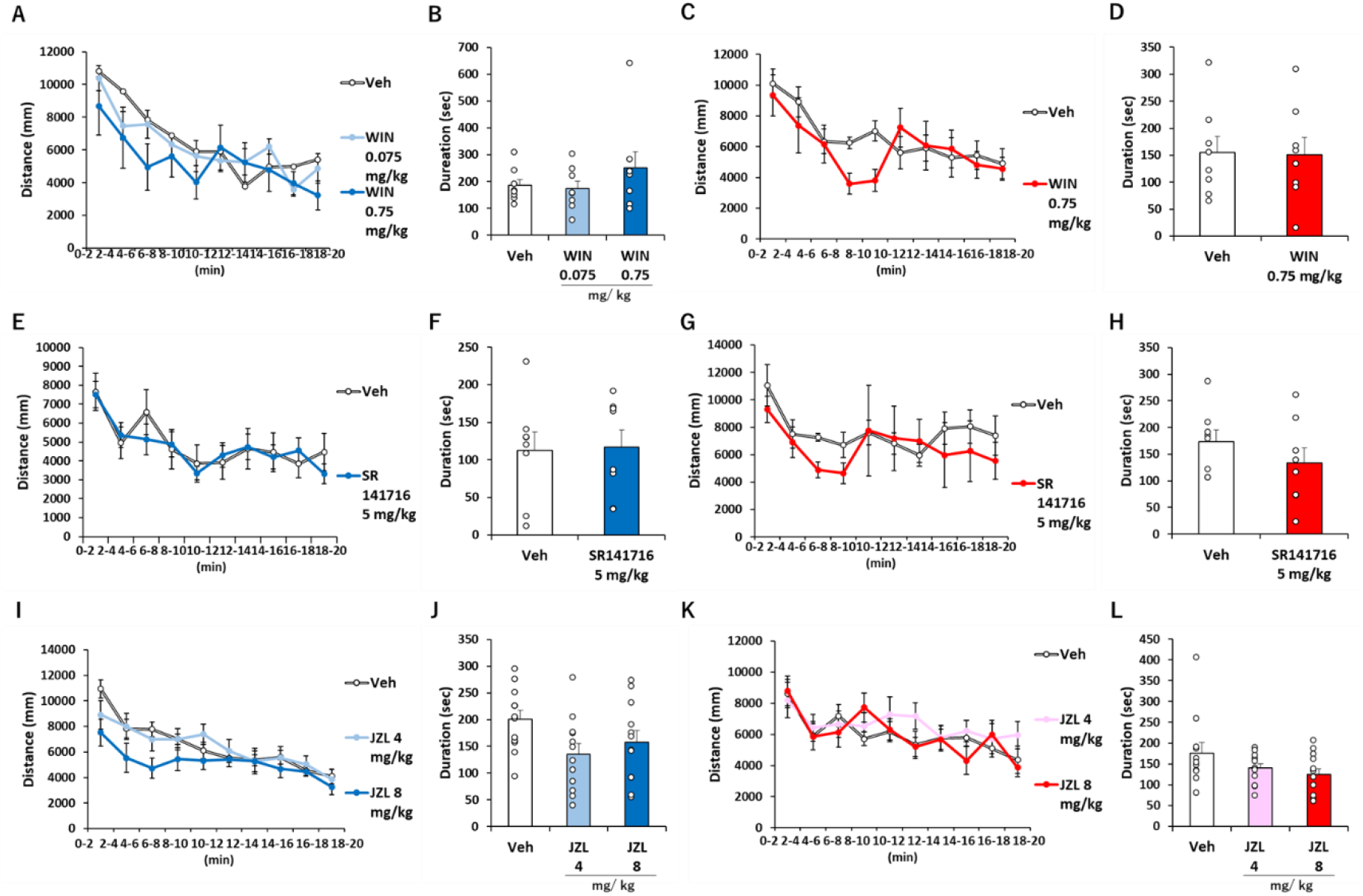
WIN, SR141716, and JZL each did not affect the total distance or the time spent in the center of the chamber. A–D: WIN (0.75 mg/kg) vs. Veh. A,C: Distance (mm) during the OFT in males (**A**) and in females (**C**). Duration (sec) of the center area in the males (**B**) and females (**D**). E–H: SR141716 (5 mg/kg) vs. Veh. Distance (mm) during the OFT in the males (**E**) and females (**G**). Duration (sec) of the center area in the males (**F**) and females (**H**). I–L: JZL (4 mg/kg and 8 mg/kg) vs. Veh. Distance (mm) during the OFT in the males (**I**) and in females (**K**). Duration (sec) of center area in the males (**J**) and in females (**L**). White circles: each animal’s data (n= 8-12/ group).

## 4 Discussion

Based on our above-described findings, we propose that systemic exogeneous stimulations of CB receptors augment the retrieval of fear memory in both sexes of mice, whereas systemic physiological eCBs do not affect this in either sex. Moreover, the increase in AEA levels does not affect the retrieval of fear memory in either sex. The increase in 2-AG levels, notably, augmented the retrieval of fear memory in only the female mice. This augmentation was regulated by the activation of CB1Rs but not CB2Rs in the females. In fear extinction, the physiological levels of CB1R activity enhance fear extinction but excess levels of CB1R activity inhibit fear extinction in both sexes.

### 4.1 No sex difference in the effect of WIN or SR141716 on the retrieval of fear memory

There was no sex difference in the effects of WIN or SR141716 on the retrieval of fear memory in this study. WIN (0.75 mg/kg) augmented the retrieval of fear memory in the males (Fig. 1B). This result is consistent with reports that the CBR agonists CP55,940 (50 μg/kg) and delta-9-tetrahydrocannabinoid (THC) (3 mg/kg) augmented the retrieval of fear memory in male mice and rats, respectively (Llorente-Berzal et al., 2015; Metna-Laurent et al., 2012). We observed herein that WIN (0.75 mg/kg) augmented the retrieval of fear memory in females to the same extent as in males (Fig. 1E). WIN (0.75 mg/kg) did not affect anxiety-like behavior in either sex (Fig. 6B,D). The effect of WIN may therefore be specific for the retrieval of fear memory rather than entirety of the negative emotion in both sexes.

In contrast, SR141716 (1 or 5 mg/kg) did not affect the retrieval of fear memory in either sex (Fig. 2B,E). Marsicano et al. (2002) reported that the retrieval of fear memory did not differ between CB1R KO mice and CB1R WT mice. It was later reported that diacylglycerol lipase (DAGL) KO male and female mice (which are deficient in 2-AG signaling) exhibited normal retrieval of fear memory comparable to that of WT mice (Cavener et al., 2018). These results suggest that systemic exogeneous stimulation of CB1Rs augments the retrieval of fear memory in both sexes, but the physiological level of systemic eCBs do not affect it in either sex.

### 4.2 Sex differences in the effect of JZL on the retrieval of fear memory

There were sex differences in the effect of JZL on the retrieval of fear memory in this study; JZL did not affect the retrieval of fear memory in the males (Fig. 3B). This result is consistent with reports that MAGL KO male mice and male mice treated with JZL (8 mg/kg), with which each treatment the 2-AG signaling was increased, did not exhibit altered retrieval of fear memory (Kishimoto et al., 2015; Llorente-Berzal et al., 2015). Unexpectedly, in the present study the treatment with JZL (8 mg/kg) augmented the retrieval of fear memory in the female mice (Fig. 3E). This augmentation was reduced by a systemic administration of SR141716 (5 mg/kg), but not by SR144528 (1 mg/kg) (Fig. 4), indicating that CB1R was involved in the augmentation. To the best of our knowledge, this study is the first to demonstrate that the increase in 2-AG levels may augment the retrieval of fear memory.

Although both WIN and 2-AG act as full agonists at CB1Rs, WIN (0.75 mg/kg) but not JZL (8 mg/kg) augmented the retrieval of fear memory in the male mice as described above.

This difference may be due to different activation levels of CB1Rs in the fear-related brain regions affected by these drugs. JZL activates CB1Rs through the increase in 2-AG levels (Long et al., 2009; Pan et al., 2009). However, 2-AG has lower affinity and less potency for CB1Rs than WIN (Presley et al., 2016). The activation of CB1Rs suppresses the release of GABA in the hippocampus (Pan et al., 2009). Interestingly, WIN reduced the amplitude of effective unitary IPSCs at dendritic connections in the hippocampus in male mice, but JZL did not (Lee et al., 2015). Moreover, CB1R-positive fiber terminals were shown to be tightly attached to parvalbumin (PV)-positive interneurons in the hippocampus that regulate the retrieval of fear memory in male mice (Sans-Dublanc et al., 2020), whereas MAGL is not expressed in these interneurons in male rats (Rivera et al., 2014). Although JZL increased the 2-AG levels by inhibiting MAGL activity (Wiskerke et al., 2012), JZL did not increase the 2-AG levels in the synaptic clefts between CB1R-positive fiber terminals and the PV-positive interneurons in the hippocampus. These results suggested that JZL less activates CB1Rs than WIN in the hippocampus in males. In addition, CB1R is expressed in the mPFC which is necessary for the retrieval of fear memory (Bisby et al., 2020; Senn et al., 2014). However, 2-AG levels were not increased after retrieval of fear memory in the medial prefrontal cortex (mPFC), although they were increased in the BLA in male mice (Marsicano et al., 2002), indicating that JZL does not increase 2-AG levels in the mPFC during the retrieval of fear memory. Taken together, these findings imply that because an intense activation of CB1Rs is required to augment the retrieval of fear memory (Metna-Laurent et al., 2012), the activation of CB1Rs in the hippocampus and mPFC by JZL may not be sufficient for increasing the retrieval of fear memory in males.

We observed that unlike the results in the males, JZL augmented the retrieval of fear memory through CB1Rs in the females. This sex difference was not caused by sex differences in functions or downstream molecules of CB1Rs, because our results demonstrated that a reduction of systemic CB1R activation did not affect the retrieval of fear memory, but an increase in CB1R activation augmented the retrieval in both sexes (Figs. 1, 2). Rather, the key may be sex differences in how much the level of 2-AG is increased by JZL. Female rats had higher DAGL mRNA levels in the frontal cortex than male rats (Marco et al., 2014), and the expression levels of MAGL in the amygdala of female rats were higher compared to those of male rats (Krebs-Kraft et al., 2010). These results suggest that the turnover of 2-AG might be faster in females than in males, and that JZL might more greatly increase the 2-AG levels in females than in males. In other words, in only females, JZL might sufficiently induce the activation of CB1Rs that is required for an increase in the retrieval of a fear memory. Further research is needed to test this possibility.

### 4.3 No sex difference in the effects of drugs targeting the CB system on fear extinction

The extent of the contribution or involvement of CB1Rs in fear extinction depends on the CB1R activation levels. For example, a low dose and a high dose of WIN (0.25 and 2.5 mg/kg) or JZL (0.5 and 8 mg/kg) facilitated and inhibited fear extinction in male rodents, respectively (Hartley et al., 2016; Llorente-Berzal et al., 2015; Morena et al., 2018; Pamplona et al., 2006). Compared to WT mice, both MAGL KO mice (which have excessively increased 2-AG signaling) and CB1R KO mice exhibit slower extinction (Kishimoto et al., 2015; Marsicano et al., 2002). The CB1R antagonist SR141716 (3 mg/kg) also inhibited fear extinction in male mice (Marsicano et al., 2002; Suzuki et al., 2004). Thus, physiological-to-moderate levels of CB1R activity enhance fear extinction, but excess levels of it inhibit fear extinction in male rodents.

We observed inhibitory effects of WIN, SR141716, and JZL on fear extinction not only in males but also in females (Fig. 1-3). Fear extinction in the proestrous phase is more fully consolidated (Milad et al., 2009). In the present study, the percentages of mice in the proestrus phase were comparable between the drug-administered groups in all extinction sessions (Suppl. Fig. S1), which suggests that the inhibitory effects of WIN, SR141716, and JZL in the female mice were not influenced by the estrous cycle. An earlier study also showed that DAGL KO mice (which are deficient in 2-AG synthesis) exhibited slower extinction than WT mice in both sexes (Cavener et al., 2018). The bell-shaped contribution of the CB1 receptors in fear extinction is thus common to both sexes.

### 4.4 No sex difference in the effect of URB597 on the retrieval and extinction of fear memory

The FAAH inhibitor URB did not affect the retrieval of fear memory in either sex of mice in this study, and this result is consistent with another recent report (Morena et al., 2021). In contrast to systemic administration, a microinjection of AEA into the left dorsolateral periaqueductal gray (dlPAG) or bilateral infralimbic cortex (IL) impaired the retrieval of fear memory in male rats (Lisboa et al., 2010; Resstel et al., 2008), and the overexpression of FAAH in the hippocampus augmented the retrieval of fear memory in male mice (Zimmermann et al., 2019). The effects of an increase in AEA on the retrieval of fear memory have thus differed between systemic administration and microinjection in male rodents. It is unclear whether such differences are observed in female rodents since no study has investigated the effects of AEA on the retrieval of fear memory in the fear-related regions in females. Such studies are merited.

We also observed that URB did not affect fear extinction in either sex (Fig. 4). This result is inconsistent with a report that URB treatment enhanced fear extinction through the activation of CB1Rs in male rats but inhibited it via an activation of transient receptor potential vanilloid 1 (Trpv1), which is a target receptor of AEA, in female rats (Morena et al., 2021). This inconsistency may be due to the differences in behavioral methods. Morena et al. used a cued fear extinction paradigm, whereas we used a contextual fear extinction paradigm. An earlier study by Morena et al. (2017) indicated that the same dose of URB did not affect contextual fear extinction in male rats, which supports the behavioral-method difference as a reason for the discrepant findings.

### 4.5 Sex differences in the duration of effects of WIN or JZL on fear extinction

As illustrated in Figure 1, Figure 3 and Supplemental Figure S4, the inhibitory effects of both WIN (0.75 mg/kg) and JZL (8 mg/kg) on FE were reduced through a 5-day extinction session in female mice, but not in male mice. WIN (0.75 mg/kg) increased the %freezing during all bins from day FE1 to day FE5 in the males, but only until bin 6 on day FE4 and only in bins 2–4 on day FE5 in the females. Moreover, when we focused on day FE5, we observed that JZL (8 mg/kg) increased the %freezing in the males but not in the females (Fig. 3).

Parks et al. (2020) reported that a reduction of the effect of CBR agonists in female mice was also produced by the antinociceptive effect of tetrahydrocannabinol (THC). When mice of both sexes were exposed to THC (10 mg/kg) for 5 consecutive days and its antinociceptive effect was investigated throughout this 5-day period, THC’s antinociceptive effect was observed for all 5 days in the male mice but its effects disappeared from day 4 in the female mice. Another study showed that after the exposure of rats to THC (30 mg/kg) for six consecutive days, the maximal [3H]SR141716A saturation binding (Bmax) decreased in both sexes, but the magnitude of the alterations was greater in the female rats (Farquhar et al., 2019). Thus, females tend to show more desensitization and/or downregulation of CB1Rs than male after sub-chronic treatment with THC.

In their study of only male mice, Nealon et al. 2019 and Schlosburg et al., 2010 reported that sub-chronic treatment with WIN and with JZL resulted in tolerance to the antinociceptive effect and desensitization, respectively. The mice were administered WIN (10 mg/kg) once-daily for nine consecutive days, and the antinociceptive effect of WIN (10 mg/kg) was evaluated on day 10: the effect of was low compared to that on day 0. Another investigation revealed that a sub-chronic increase in 2-AG levels by the administration of JZL (40 mg/kg) to male mice for 6 days resulted in CB1R desensitization and/or downregulation in the cingulate cortex, hippocampus, somatosensory cortex, and periaqueductal gray (Schlosburg et al., 2010). Although it is unclear whether there are sex differences in the reduction of effects of WIN on antinociception and CB1R desensitization and/or downregulation by 2-AG (Nealon et al., 2019; Schlosburg et al., 2010), a possible explanation for the female-dominant reduction of the effects of the pharmacological activation of CB1Rs on fear extinction in our present study might be that a sub-chronic activation of CB1Rs induced by WIN or JZL may induce more desensitization and/or downregulation of CB1Rs in the fear-related brain regions of females compared to those of males.

### 4.6 Inconsistency with previous studies using WIN or JZL

The effects of WIN and JZL on the retrieval and extinction of fear memory in the present study are inconsistent with those of many reports that the activation of CB1Rs by WIN, JZL, or other CBR agonists reduces the retrieval of fear memory and/or enhances fear extinction in male rodents (Atsak et al., 2012; Bisby et al., 2020; Bitencourt et al., 2008; CG Reich et al., 2008; Ghasemi et al., 2017; Korem et al., 2017; Kuhnert et al., 2013; Lin et al., 2009; Lisboa et al., 2015; Morena et al., 2017; Pamplona et al., 2006, 2008; Sachser et al., 2015; Segev and Akirav, 2011; Simone et al., 2015). This inconsistency may be caused by differences in experimental methods. The role of the CB1Rs in the regulation of fear depends on the activation levels of CB1Rs (Metna-Laurent et al., 2012; Pamplona et al., 2006). Moreover, the activation levels of CB1Rs for reducing a fear memory are influenced by factors including species, age, drugs, and foot shock intensity (Bisby et al., 2020; Haller et al., 2007; Jacob et al., 2012).

Of those factors, the effect of the species is likely to affect the inconsistency the most. Our present findings are consistent with those of studies using mice (Llorente-Berzal et al., 2015) but inconsistent with those of studies using rats (Atsak et al., 2012; Bisby et al., 2020; Bitencourt et al., 2008; CG Reich et al., 2008; Ghasemi et al., 2017; Korem et al., 2017; Kuhnert et al., 2013; Lin et al., 2009; Lisboa et al., 2015; Pamplona et al., 2006, 2008; Sachser et al., 2015; Segev and Akirav, 2011; Simone et al., 2015). The hippocampal IPSCs in mice showed larger sensitivity to WIN compared to those in rats (Haller et al., 2007). The activation of inhibitory neurons in the hippocampus is required to reduce the retrieval of a fear memory (Sans-Dublanc et al., 2020), and because the activation of CB1Rs suppresses the release of GABA in the hippocampus(Pan et al., 2009), the excitability of hippocampal neurons in mice might be more greatly increased by the activation of CB1Rs compared to those in rats.

## 5. Conclusion

JZL (an inhibitor of the 2-AG MAGL) augmented the retrieval of a fear memory in this study via the activation of CB1Rs, but not CB2Rs, in only female mice. The endocannabinoid 2-AG regulates the release of various neurotransmitters such as glutamate, GABA, acetylcholine, and dopamine (Griebel et al., 2015; Oleson et al., 2012; Pan et al., 2009).These neurotransmitters are related to many physiological functions, behaviors, and psychiatric disorders (Changeux, 2010; Chiu et al., 2005; Gu, 2002; Javitt, 2004; Perez de la Mora et al., 2020). Our present findings suggest the importance of investigating the effects of the augmentation of 2-AG on various physiological functions, behaviors, and psychiatric disorders by using females, even if it is known that 2-AG does not influence these in males.

## Author contributions

Ikumi Mizuno: Investigation, Visualization, Writing - original draft, Writing – review & editing. Shingo Matsuda: Conceptualization, Formal analysis, Supervision, Funding acquisition, Project administration, Methodology, Writing - original draft, Data curation, Writing -review & editing. Suguru Tohyama: Writing -review & editing. Akihiro Mizutani: Writing - original draft, Writing -review & editing, Funding acquisition.

## Acknowledgements

Shingo M. conceived this research; Ikumi M. and Shingo M. designed this research and performed the experiments; Shingo M. analyzed the data; All authors interpreted experimental results, drafted the manuscript, and commented on the manuscript. We would like to thank Yuri Furuya and Yoshimi Ogawa for assisting with performing the experiments.

