## Supplement for "The role of the cannabinoid system in fear memory and extinction in male and female mice"

**A**

**Veh**

| Animal ID | Day 1 | Day 2 | Day 3 | Day 4 | Day 5 |
|---|---|---|---|---|---|
| f1 | e | e | d | d | d |
| f2 | d | e | d | d | d |
| f3 | d | d | d | d | p |
| f4 | d | d | d | p | e |
| f5 | e | e | e | d | d |
| f6 | e | d | d | d | p |
| f7 | d | d | d | p | e |
| f8 | d | d | d | e | e |
| f9 | e | d | d | p | e |
| f10 | d | d | d | d | d |
| f11 | d | d | d | d | d |
| f12 | d | d | d | p | e |
| f13 | e | d | d | d | d |
| P (%) | 0 | 0 | 0 | 30.77 | 15.385 |

**WIN 0.075 mg/kg**

| Animal ID | Day 1 | Day 2 | Day 3 | Day 4 | Day 5 |
|---|---|---|---|---|---|
| f1 | d | d | d | d | d |
| f2 | d | d | d | e | e |
| f3 | e | e | d | d | d |
| f4 | d | d | p | e | e |
| f5 | d | d | d | d | p |
| f6 | d | d | d | d | e |
| f7 | d | d | d | d | d |
| P (%) | 0 | 0 | 14.29 | 0 | 14.286 |

**WIN 0.75 mg/kg**

| Animal ID | Day 1 | Day 2 | Day 3 | Day 4 | Day 5 |
|---|---|---|---|---|---|
| f1 | d | p | e | d | d |
| f2 | e | d | d | d | p |
| f3 | d | d | d | p | e |
| f4 | e | e | e | d | d |
| f5 | e | d | d | d | d |
| f6 | e | d | d | d | d |
| f7 | e | e | d | d | d |
| f8 | p | d | d | d | d |
| f9 | d | e | d | d | d |
| f10 | pe | d | d | d | d |
| f11 | d | p | e | e | e |
| f12 | p | e | d | d | d |
| f13 | d | d | d | dp | e |
| P (%) | 15.385 | 15.38 | 0 | 7.692 | 7.6923 |

**B**

**Veh**

| Animal ID | Day 1 | Day 2 | Day 3 | Day 4 | Day 5 |
|---|---|---|---|---|---|
| f1 | d | d | p | e | e |
| f2 | e | d | d | d | p |
| f3 | d | d | p | e | e |
| f4 | p | e | e | d | d |
| f5 | d | d | p | e | e |
| f6 | d | d | d | d | d |
| f7 | d | d | d | d | p |
| f8 | d | d | d | e | e |
| f9 | d | d | d | d | d |
| f10 | d | d | d | d | d |
| f11 | e | e | d | d | p |
| P (%) | 9.09 | 0 | 27.27 | 0 | 27.27 |

**SR141716 1 mg/kg**

| Animal ID | Day 1 | Day 2 | Day 3 | Day 4 | Day 5 |
|---|---|---|---|---|---|
| f1 | d | d | d | d | d |
| f2 | p | e | e | e | d |
| f3 | e | e | e | d | d |
| f4 | e | e | d | d | d |
| f5 | d | d | d | p | e |
| f6 | e | d | d | d | p |
| f7 | d | d | d | d | p |
| f8 | d | d | d | p | e |
| f9 | d | p | e | d | d |
| f10 | d | d | d | p | e |
| f11 | d | d | d | d | d |
| P (%) | 9.09 | 9.09 | 0 | 27.27 | 18.18 |

**SR141716 5 mg/kg**

| Animal ID | Day 1 | Day 2 | Day 3 | Day 4 | Day 5 |
|---|---|---|---|---|---|
| f1 | d | d | d | d | d |
| f2 | d | d | p | e | e |
| f3 | ed | d | d | d | d |
| f4 | d | d | d | d | e |
| f5 | d | d | d | d | e |
| f6 | p | e | e | d | d |
| f7 | e | d | d | d | e |
| f8 | d | d | p | e | e |
| f9 | e | d | d | d | d |
| f10 | e | d | d | d | d |
| f11 | e | d | d | d | e |
| P (%) | 9.09 | 0 | 18.18 | 0 | 0 |

**C**

**Veh**

| Animal ID | Day 1 | Day 2 | Day 3 | Day 4 | Day 5 |
|---|---|---|---|---|---|
| f1 | d | d | d | d | d |
| f2 | e | d | d | d | p |
| f3 | d | d | d | d | d |
| f4 | d | p | d | d | d |
| f5 | e | e | e | d | d |
| f6 | e | e | d | d | d |
| f7 | d | e | e | e | e |
| f8 | p | e | e | d | d |
| f9 | d | d | d | d | d |
| f10 | d | d | d | d | d |
| f11 | d | d | d | p | e |
| f12 | d | d | d | d | d |
| f13 | d | d | d | d | d |
| f14 | d | d | d | d | d |
| f15 | d | d | d | d | d |
| f16 | d | d | d | e | e |
| f17 | e | d | d | d | p |
| P (%) | 5.88 | 5.88 | 0 | 5.88 | 11.76 |

**JZL 4 mg/kg**

| Animal ID | Day 1 | Day 2 | Day 3 | Day 4 | Day 5 |
|---|---|---|---|---|---|
| f1 | p | e | e | e | d |
| f2 | d | d | d | d | d |
| f3 | e | e | d | d | d |
| f4 | d | d | d | d | d |
| f5 | e | d | d | d | d |
| f6 | d | e | d | d | d |
| f7 | e | e | d | d | d |
| f8 | d | dp | e | d | d |
| f9 | p | e | d | d | p |
| f10 | d | d | d | d | p |
| f11 | e | e | d | d | d |
| f12 | e | d | d | d | p |
| f13 | p | e | d | d | p |
| f14 | d | d | d | d | d |
| f15 | e | d | d | d | d |
| f16 | d | d | d | d | d |
| f17 | e | e | d | d | d |
| f18 | d | d | d | d | d |
| f19 | e | e | e | e | d |
| P (%) | 17.65 | 0 | 0 | 0 | 23.53 |

**JZL 8 mg/kg**

| Animal ID | Day 1 | Day 2 | Day 3 | Day 4 | Day 5 |
|---|---|---|---|---|---|
| f1 | d | d | d | d | d |
| f2 | d | d | d | d | d |
| f3 | d | d | d | d | d |
| f4 | d | d | d | d | d |
| f5 | e | d | d | d | d |
| f6 | d | d | d | d | d |
| f7 | d | d | d | dp | e |
| f8 | d | d | e | e | ed |
| f9 | e | e | e | d | d |
| f10 | e | d | e | d | d |
| f11 | e | d | d | d | d |
| f12 | d | d | d | d | d |
| f13 | d | d | d | d | d |
| f14 | e | d | d | d | p |
| f15 | e | d | d | d | d |
| f16 | d | d | d | d | p |
| f17 | e | e | d | d | d |
| f18 | d | d | d | d | p |
| P (%) | 0 | 0 | 0 | 0 | 17.65 |

**D**

**Veh**

| Animal ID | Day 1 | Day 2 | Day 3 | Day 4 | Day 5 |
|---|---|---|---|---|---|
| f1 | e | e | d | d | d |
| f2 | d | d | d | d | p |
| f3 | e | d | d | d | e |
| f4 | e | d | d | d | d |
| f5 | e | e | d | d | d |
| f6 | d | d | d | p | e |
| f7 | d | d | d | d | dp |
| f8 | d | p | e | d | d |
| f9 | e | e | e | e | d |
| f10 | e | d | d | d | p |
| f11 | d | d | p | e | e |
| f12 | e | e | d | d | d |
| P (%) | 0 | 8.33 | 8.33 | 8.33 | 16.67 |

**URB 0.3 mg/kg**

| Animal ID | Day 1 | Day 2 | Day 3 | Day 4 | Day 5 |
|---|---|---|---|---|---|
| f1 | d | d | e | e | d |
| f2 | d | p | e | d | d |
| f3 | p | e | e | e | d |
| f4 | e | d | d | d | d |
| f5 | p | e | e | d | d |
| f6 | e | e | d | d | d |
| f7 | d | d | p | e | e |
| f8 | p | e | e | d | d |
| f9 | p | e | e | d | d |
| f10 | d | d | d | d | d |
| f11 | ed | d | d | d | pe |
| f12 | e | d | d | d | p |
| P (%) | 33.33 | 8.33 | 8.33 | 0 | 8.33 |

**URB 1 mg/kg**

| Animal ID | Day 1 | Day 2 | Day 3 | Day 4 | Day 5 |
|---|---|---|---|---|---|
| f1 | d | d | d | d | d |
| f2 | d | d | e | d | d |
| f3 | p | e | e | d | d |
| f4 | p | e | e | d | d |
| f5 | d | d | d | e | e |
| f6 | d | d | e | ed | d |
| f7 | e | d | d | d | p |
| f8 | d | d | d | d | d |
| f9 | d | d | p | e | e |
| f10 | d | d | d | d | p |
| f11 | d | d | d | d | d |
| f12 | e | d | d | d | d |
| P (%) | 16.67 | 0 | 8.33 | 0 | 16.67 |

**URB 3 mg/kg**

| Animal ID | Day 1 | Day 2 | Day 3 | Day 4 | Day 5 |
|---|---|---|---|---|---|
| f1 | d | d | d | d | d |
| f2 | d | d | d | e | d |
| f3 | d | d | ed | e | d |
| f4 | d | d | d | p | e |
| f5 | e | e | d | d | d |
| f6 | d | d | d | d | d |
| f7 | d | d | d | d | p |
| f8 | d | d | d | d | dp |
| f9 | d | d | d | d | d |
| f10 | d | d | d | d | d |
| f11 | d | d | e | e | e |
| f12 | d | d | d | d | p |
| P (%) | 0 | 0 | 0 | 8.33 | 16.67 |

**Suppl. Fig. S1.** Estrous cycle phase of the female mice on each FE day. The estrous cycle phase in the experiment investigating the effects of WIN (**A**), SR141716 (**B**), JZL (**C**) and URB (**D**) on the retrieval or extinction of fear memory. D: diestrous phase, DP: transitional phase from D to P, E: estrous phase, P (%): The percentage of proestrous phase, P: proestrous phase, PE: transitional phase from P to E, ED: transitional phase from E to D.

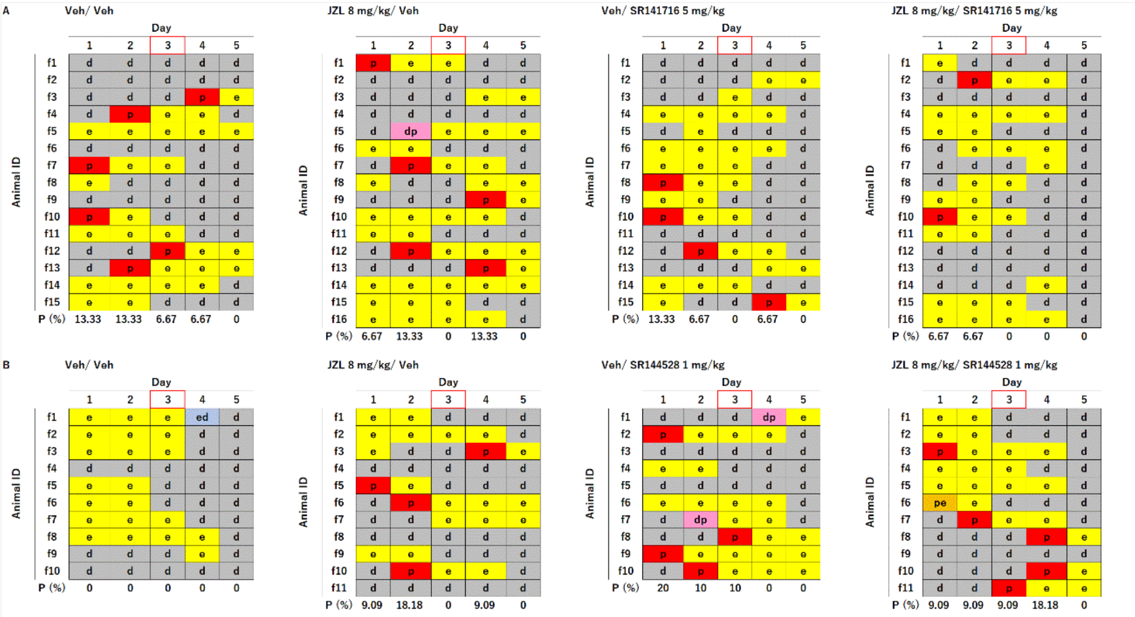


**Suppl. Fig. S2.** The estrous cycle phase on the day of the TEST. The estrous cycle phase in the experiment investigating the effects of JZL and SR141716 (**A**), JZL and SR144528 (**B**) on the retrieval of fear memory. The *red frame* represents the TEST day. Abbreviations are explained in the Suppl. Fig. S1 legend.

**A** Veh

| Animal ID | Day 1 | Day 2 | Day 3 | Day 4 | Day 5 |
|---|---|---|---|---|---|
| f1 | p | e | e | d | d |
| f2 | d | d | p | e | d |
| f3 | d | d | p | e | e |
| f4 | d | d | e | d | d |
| f5 | p | e | e | d | d |
| f6 | p | e | e | e | e |
| f7 | e | d | d | | e |
| f8 | p | e | e | | p |
| P (%) | 50 | 0 | 25 | 0 | 12.50 |

WIN 0.75 mg/kg

| Animal ID | Day 1 | Day 2 | Day 3 | Day 4 | Day 5 |
|---|---|---|---|---|---|
| f1 | d | d | d | p | e |
| f2 | e | e | e | d | d |
| f3 | e | e | e | e | e |
| f4 | e | e | e | d | d |
| f5 | p | e | e | d | d |
| f6 | e | e | e | e | d |
| f7 | p | e | e | d | d |
| f8 | p | e | e | d | d |
| P (%) | 37.5 | 0 | 0 | 12.5 | 0 |

**B** Veh

| Animal ID | Day 1 | Day 2 | Day 3 | Day 4 | Day 5 |
|---|---|---|---|---|---|
| f1 | d | d | d | p | e |
| f2 | e | e | pe | e | e |
| f3 | e | e | e | d | d |
| f4 | d | d | d | d | p |
| f5 | d | d | d | p | e |
| f6 | p | e | e | e | d |
| f7 | p | e | e | d | d |
| f8 | d | d | d | d | d |
| P (%) | 25 | 0 | 0 | 25 | 12.5 |

SR141716 5 mg/kg

| Animal ID | Day 1 | Day 2 | Day 3 | Day 4 | Day 5 |
|---|---|---|---|---|---|
| f1 | e | d | d | d | d |
| f2 | e | d | d | d | d |
| f3 | d | e | e | e | d |
| f4 | d | d | e | e | e |
| f5 | e | d | d | p | e |
| f6 | d | d | e | e | e |
| f7 | p | e | e | d | d |
| f8 | e | e | d | d | p |
| P (%) | 12.5 | 0 | 0 | 12.5 | 12.5 |

**C** Veh

| Animal ID | Day 1 | Day 2 | Day 3 | Day 4 | Day 5 |
|---|---|---|---|---|---|
| f1 | e | e | e | d | d |
| f2 | d | d | e | e | e |
| f3 | e | e | d | d | e |
| f4 | d | d | d | e | e |
| f5 | e | e | e | e | e |
| f6 | p | e | e | d | d |
| f7 | e | e | e | d | d |
| f8 | d | d | d | e | e |
| f9 | d | d | d | d | d |
| f10 | d | d | d | p | e |
| f11 | d | d | d | d | d |
| f12 | d | p | e | e | e |
| P (%) | 8.33 | 8.33 | 0.00 | 8.33 | 0.00 |

JZL 4 mg/kg

| Animal ID | Day 1 | Day 2 | Day 3 | Day 4 | Day 5 |
|---|---|---|---|---|---|
| f1 | d | d | p | e | ed |
| f2 | d | d | e | e | e |
| f3 | d | d | d | d | e |
| f4 | d | d | d | e | e |
| f5 | d | e | d | d | d |
| f6 | d | e | e | e | e |
| f7 | e | e | e | d | d |
| f8 | d | d | d | d | e |
| f9 | d | d | d | dp | e |
| f10 | e | e | e | e | e |
| f11 | dp | e | e | e | e |
| f12 | p | e | e | e | e |
| P (%) | 8.33 | 0.00 | 8.33 | 0.00 | 0.00 |

JZL 8 mg/kg

| Animal ID | Day 1 | Day 2 | Day 3 | Day 4 | Day 5 |
|---|---|---|---|---|---|
| f1 | e | ed | d | d | e |
| f2 | e | e | d | d | e |
| f3 | d | d | d | e | e |
| f4 | p | e | e | d | d |
| f5 | p | e | e | e | e |
| f6 | e | e | d | d | e |
| f7 | e | e | e | d | d |
| f8 | d | d | p | e | e |
| f9 | d | e | e | e | e |
| f10 | d | p | e | e | e |
| f11 | p | e | e | e | e |
| f12 | p | e | e | d | d |
| P (%) | 33.33 | 8.33 | 8.33 | 0.00 | 0.00 |

**Suppl. Fig. S3.** The estrous cycle phase on the day that the OFT was conducted. The estrous cycle phase in the experiment investigating the effect of WIN (**A**), SR141716 (**B**), and JZL (**C**). *Red frame:* the OFT day. Abbreviations are explained in the Suppl. Fig. S1 legend.

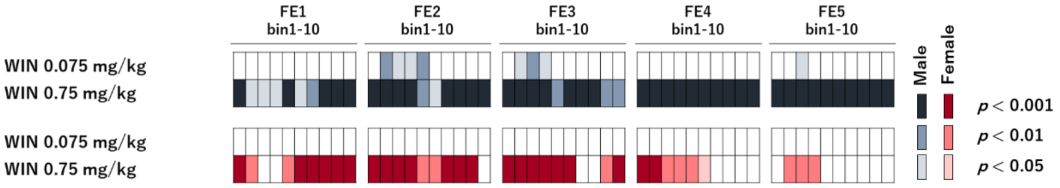


**Suppl. Fig. S4.** The statistical results of the post-hoc analyses showing the interaction of bin × group × FE day in the experiment using WIN. *Colored oblong:* significant differences between Veh and WIN.

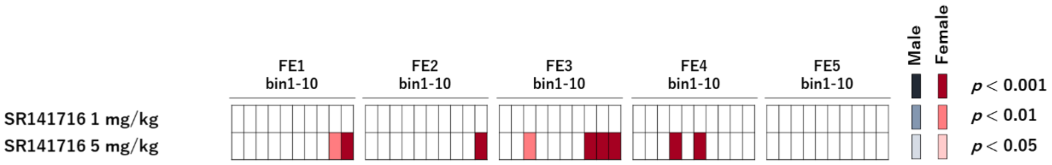


**Suppl. Fig. S5.** The statistical results of the post-hoc analyses showing the interaction of bin × group × FE day in the experiment using SR141716. *Colored oblong:* significant differences between Veh and SR141716.

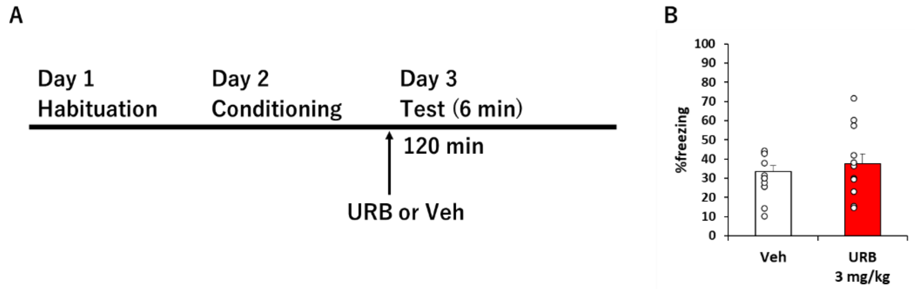


**Suppl. Fig. S6.** URB did not affect the retrieval of fear memory in the female mice. **A:** The procedure of experiment 5. The *arrow* indicates the time point of drug injection that was performed 120 min before the TEST. **B:** The percentage of freezing during the TEST in females. *White circles:* each animal's data (n=12/group). There was no significant difference in %freezing among the groups: $t$ (22) = −1.219, $p$= 0.236.

Suppl. Fig. S4

**Supplementary Table S1 The results of one way-ANOVA and Student's t-test in OFT**

| Sex | Drug | Total distance | | Center time |
|---|---|---|---|---|
| | | **Main effect Group** | **Interaction bin × Group** | **Main effect Group or Student's t-test** |
| Male | WIN | $F(2, 21) = 1.357$ $p= 0.28$ | $F(9.492, 99.669) = 1.017$ $p= 0.433$ | $F(2, 22) = 1.076$ $p= 0.359$ |
| | SR141716 | $F(1, 14) = 0.30$ $p= 0.864$ | $F(4.385, 61.393) = 0.980$ $p= 0.430$ | $t$ (14) = -0.131 $p= 0.897$ |
| | JZL | $F(2, 33) = 1.857$ $p= 0.172$ | $F(10.769, 177.684) = 1.492$ $p= 0.140$ | $F(2, 34) = 2.831$ $p= 0.073$ |
| Female | WIN | $F(1, 14) = 0.922$ $p= 0.353$ | $F(4.407, 61.696) = 1.081$ $p= 0.377$ | $t$ (14) = 0.078 $p= 0.939$ |
| | SR141716 | $F(1, 14) = 0.444$ $p= 0.516$ | $F(2.806, 39.289) = 0.687$ $p= 0.556$ | $t$ (14) = 1.146 $p= 0.271$ |
| | JZL | $F(2, 33) = 0.500$ $p= 0.611$ | $F(11.223, 185.175) = 0.956$ $p= 0.490$ | $F(2, 34) = 2.295$ $p= 0.117$ |
